# Arctic greening drives changes in the diet and gut microbiome of a resident herbivore with consequences for fitness

**DOI:** 10.1101/2024.02.27.582426

**Authors:** Stefaniya Kamenova, Steve D. Albon, Leif Egil Loe, R. Justin Irvine, Rolf Langvatn, Galina Gusarova, Eric Jacques de Muinck, Pål Trosvik

## Abstract

Rapid climate warming is “greening” the tundra due to higher plant productivity, particularly of deciduous shrubs and grasses, potentially changing the nutritional base for herbivores. However, the consequences for herbivores are unclear. Although the gut microbiome is an essential factor, mediating the capacity to adapt to dietary change, few studies have investigated the effect of annual changes in the diet-gut microbiome nexus on fitness traits. In a model system, the Svalbard reindeer, a species experiencing the greatest rate of climate warming on Earth, we investigate how climate-induced changes in diet and the gut microbiome impact reindeer body mass. Using high-resolution DNA metabarcoding, we quantified diet and microbiome composition in rumen samples from 97 animals culled in October from 1998 to 2004 in four different valleys. Overall diet diversity and grass consumption were significantly higher following warmer summers, while intake of the dwarf shrub *Salix polaris* increased with the maximum normalised difference vegetation index (NDVI). The proportion of *Salix* had a significant positive effect on autumn body mass, which in turn increases ovulation rates and overwinter survival. There was a significant positive effect of *Salix* on the microbiome diversity, while between-year variation in Bacteroidota, the most common bacteria phylum, was also significantly, positively related to maximum NDVI. However, a structural equation model revealed that the direct “path” effect of the proportion of *Salix* in the diet on reindeer body mass was stronger than the indirect path effect, mediated through the gut microbiome. Our results suggest a positive impact of climate-driven Arctic greening on Svalbard reindeer fitness, operating through a change in diet composition, partially mediated by the changes in the gut microbiome. These findings advance our understanding of the mechanistic underpinnings of Arctic warming on herbivore populations and contribute to the debate about the importance of the diet−gut microbiome nexus in facilitating species resilience in the face of climate change.

## Introduction

The Arctic tundra is warming fast, with temperatures rising almost four times more rapidly than the global average (Rantanen et al. 2022). Such a dramatic change is already disrupting the ways species interact with each other, with consequences for tundra ecosystem functioning (Post et al. 2009). The warmer and longer plant growing seasons due to climate warming across the circumpolar Arctic has led to widespread increases in plant productivity (Myers-Smith et al. 2011; Forbes et al. 2017; Berner et al. 2020). Many of the plant species benefiting from the warming climate, including graminoids (grasses, sedges) and deciduous shrubs (e.g., birch and *Salix* dwarf shrubs) are the preferred forage for keystone large herbivores, such as caribou/reindeer and muskox (Bråthen & Oksanen 2001; Bjørkvoll et al. 2009; Larter & Nagy 2004; Thing et al. 1987). While the quantity of forage plants is generally increasing, short-term warming experiments suggest plant quality (digestibility and nutrient content) could decline (Zamin et al. 2017), though there may be some acclimation (Leffler et al. 2022). Hence, how these herbivores benefit from Arctic greening, if at all, is unclear. So, although many reindeer and caribou have undergone marked declines in population size (Vors & Boyce 2009; Fauchald et al. 2017), mainly due to the combined effect of anthropogenic pressure and extreme weather events. On a more local scale an earlier vegetation green-up and higher plant productivity can increase calf autumn body mass and female reproductive success (for example, reindeer in Northern Norway, Tveraa et al. 2013). Similarly, higher plant productivity in warm summers (Van der Wal & Stien 2014) increases autumn body mass, leading to higher ovulation rates in Svalbard reindeer (Albon et al, 2017). Interestingly, while muskox population growth in West Greenland also benefited from earlier green-up outweighing the impacts of winter weather, this was not the case in a sympatric caribou population where numbers declined because spring green-up seemed to have little impact on calf production compared to winter weather (Eikelenboom et al. 2021). Thus, the effects of Arctic greening on large mammalian herbivores might be both species- and context-dependent, raising the question about the specific niche requirements and mechanisms underpinning the direction of impacts.

Since large ruminant herbivores are particularly dependent on their foregut microbiomes for harvesting energy from the plants they ingest (Leser & Mølbak 2009; Shabat et al. 2016), it would seem prudent to also study the response of the gut microbiome to climate-induced changes in diet. Typically, the gut microbiome is a highly plastic, rapidly evolving system (Candela et al. 2012). In fact, for species with long generation times (i.e., little capacity for evolutionary adaptation) gut microbiome plasticity could be a key mechanism enabling rapid ecological adaptation to changes in the plane of nutrition (cf Alberdi et al. 2016; Clayton et al. 2016). Nevertheless, only now are we starting to comprehend the links and the feedbacks between foraging, the gut microbiome and fitness in wildlife, and the empirical validations are still limited. For example, recent findings from the giant panda show that changes in diet composition and quality could elicit adaptive response of the gut microbiome, with a direct effect on the maintenance of body mass and condition (Huang et al. 2022). However, so far, long-term studies on the diet-gut microbiome nexus have been restricted to humans, mainly showing positive long-lasting relationship between diet quality and gut microbiome diversity and functioning (Ma et al. 2022; Yu et al. 2021; Petrone et al. 2023). On the other hand, some studies have shown that the gut microbiome can also be resilient to changes in diet (Fragiadakis et al. 2020), with bacterial strains being able to persist for decades within the human gut (Faith et al. 2013). With the occurrence of such antagonistic processes and the limited set of studies to date, makes identifying the mechanisms driving the impact of Arctic greening on large herbivores a challenge.

Testing hypotheses about the mechanisms by which climate-driven changes in plant productivity impact on herbivore fitness requires long-term data on diet selection, but unfortunately these remain rare. Here, in a long-term study of Svalbard reindeer we address the question how climate-driven changes in plant productivity (Van der Wal & Stien 2014) influence the autumn diet and rumen microbiome, and the impact on individual performance, underpinning its sustained population growth (Le Moullec et al. 2019). For this, we use a set of reindeer rumen samples and body mass data, collected in late October over seven years. Using high-resolution DNA metabarcoding to quantify both diet and microbiome composition, we document annual variation in the diet and microbiota. Given the increasing plant productivity as summers warmed on Svalbard (Van der Wal & Stien 2014), we test the hypothesis that the proportion of preferred forage plants (graminoids and dwarf shrubs) in the autumn diet increases in warmer summers. Also, we predict a significant positive effect of these preferred plant functional groups on body mass, a key fitness trait influencing ovulation rates (Albon et al. 2017). Finally, we expect a significant correlation between diet and gut microbiome composition, as well as an independent positive gut microbiome-driven effect on the reindeer body mass in line with the tenets of the holobiont theory (Rosenberg & Zilber-Rosenberg 2018; Simon et al. 2019).

## Material and Methods

### Study area

Svalbard reindeer were sampled between 1998 and 2004 from two main areas in Svalbard - the Colesdalen and Semmeldalen valleys in Nordenskiöld land (77°50’ – 78°20’N, 15°00’ – 17°30’E), and the Sassendalen and Eskerdalen valleys in Sassen-Bünsow Land National Park (78°23′N, 17°15′E), Figure S1. The generally wide, U-shaped valleys are mostly vegetated up to about 250 m altitude, although above-ground live vascular plant biomass averages only c. 35 g m^2^ (annual range 23 – 46 g m^2^, Van der Wal & Stien 2014). In the drier *Luzula* heath and ridge habitats (c. 40% of the area) dwarf shrub species, particularly *Salix polaris*, account for the 61% and 78%, respectively, of the above ground biomass in early August. On the less well drained shallow slopes and marsh areas, grasses, sedges, and rushes are more dominant (Van der Wal & Stien 2014). The period 1998 – 2004 was an early phase of warming on Svalbard with higher plant productivity (Van der Wal & Stien 2014) and commensurate increase in autumn body mass of reindeer (Albon et al. 2017). During the study, the estimated reindeer population size in the Colesdalen/Semmeldalen (see Lee et al. 2015) varied between 948 (898 – 1006 ±CI) and 1339 (1251 – 1440 ±CI) individuals (estimates include lower Reindalen, Albon et al. 2017), while the Sassendalen/Eskerdalen population counts increased from ∼750 to ∼1250 (Hansen et al. 2019). Populations from the Colesdalen/Semmeldalen and the Sassendalen/Eskerdalen areas are effectively isolated from each other, with little interchange revealed by genetic analysis (Côté et al. 2002).

### Sample collection

Reindeer were culled in October (between 19^th^ and 27^th^) and body mass recorded before evisceration. From the rumen between 300 – 500 mL of content was collected, sampling from multiple locations within the rumen. All samples were stored in the same conditions until analysis – i.e., frozen at −20°C. Individuals were aged by counting rings in the cementum of the first incisor (Reimers & Nordby 1968) and lactation status (lactating or not lactating) was based on the presence of milk in the udder (Albon et al. 2017).

### Environmental variables measurements

Daily meteorological measurements were recorded at Svalbard Airport, Longyearbyen (Station SN99840, Norwegian Climate Services: https://seklima.met.no/observations) c. 40 km from the most distant valleys. Mean July temperature for the seven years of this study was 6.93°C ±0.8SD but showed no temporal trend (r = 0.052, P > 0.9). Nonetheless, across the seven years there was a significant increase in the annual maximum remote-sensed normalised difference vegetation index (NDVI, r = 0.775, P = 0.048), an indicator of greening widely accepted as a correlate of plant productivity, in particular sensitive to the expansion of shrub species (Jespersen et al. 2023). Maximum NDVI was calculated by generating Normalized Difference Vegetation Indices (NDVI3g) from the Global Inventory Modelling and Mapping Studies (GIMMS) dataset of AVHRR images at the National Centre for Climate Research, Boulder, Colorado, USA. NDVI values range from +1.0 to -1.0. Areas of snow, rock, or sand usually show very low NDVI values (0.1 or less). Sparse vegetation such as shrub tundra result in moderate NDVI values (0.2 to 0.5). In Svalbard NDVI values peak in late July/early August (Arnold et al. 2018) coinciding with peak biomass (Karlsen et al. 2018). The maximum NDVI was estimated from six 15-day composites from late July to early September each year using pixels within our study area. These annual values are well correlated (r = 0.993, N = 7, P<0.001) with those for a larger area around Istfjordan (Vickers et al. 2016).

### DNA metabarcoding diet analysis

Approximately 15 g of wet rumen material was homogenised in liquid nitrogen. DNA was extracted from 100 mg of the resulting ground rumen powder using the NucleoSpin Plant II (Macherey-Nagel, Germany) and following manufacturer’s instructions. Blank extractions (ultra-pure Milli-Q water instead of DNA) were included for every 24 samples. We targeted the plant part of the reindeer diet by using the *Sper01* primers amplifying the chloroplast trn*L* P6 loop in seed plants (Spermatophyta) (Taberlet et al. 2007). This is a commonly used DNA metabarcoding plant marker with extensive DNA reference database for the boreal and arctic regions (ArctBorBryo, Voldstad et al. 2020). PCR amplifications were carried out in a final volume of 15 μL, using the AmpliTaq Gold 360 PCR Master Mix (Thermo Fisher Scientific, USA), 2 μL of DNA extract as template, 0.4 μl/15 ml of BSA and 0.5 μM of each primer. The PCR mixture was denatured at 95°C for 10 min, followed by 35 cycles of 30 s at 95°C and 30 s at 52°C, and an elongation step for 7 min at 72°C. A 8 – 9-nt sequence tags were added on the 5’ end of each forward and reverse primer, resulting in a unique tag combination for each PCR product in order to allow the assignment of sequence reads for the relevant sample. Each PCR reaction was carried out in triplicate and PCR negative controls (ultra-pure Milli-Q water instead of DNA) were systematically included to monitor for contaminations. We also included one positive control for each 95 PCR replicates. Positive controls consisted of an artificially assembled mock community containing a mixture of varying proportions of six unique synthetic DNA stretches with varying GC content, homopolymers and sequence length (https://github.com/pheintzman/metabarcoding, Table S1). A subset of PCR products was selected for the visual inspection of the amplified DNA using 1.5% gel electrophoresis. PCR products were pooled and purified using QIAquick PCR Purification Kit (Qiagen, Germany). A Qubit 2.0 fluorometer and the dsDNA HS Assay kit (Invitrogen, Life Technologies, USA), and pooled again prior to library preparation and sequencing. Libraries were prepared using the KAPA HyperPlus kit (Kapa Biosystems, USA), and sequenced on a HiSeq 4000 machine (Illumina, USA) following manufacturer’s instructions at the Norwegian Sequencing Centre (https://www.sequencing.uio.no). A total of 150 nucleotides were sequenced on each extremity of the DNA fragments.

### Bioinformatic analyses

Sequences were analysed using the OBITools pipeline (Boyer et al. 2016). The direct and reverse reads were aligned and merged using the *illuminapairedend* command by considering the quality of the sequence data during the alignment and the consensus computation. Only alignments with scores >50 were kept for further analyses. Primers and tags were identified using the *ngsfilter* command. Only sequences with a perfect match on tags and a maximum of two errors on primers were retained for further analyses. Primers and tags were cut off at this step. Strictly identical sequences were clustered together using the *obiuniq* command, while keeping the information about their distribution among samples. All sequences shorter than 10 bp and/or occurring at ≤10 reads were excluded. Sequences corresponding to PCR and/or sequencing errors were labelled to be removed using *obiclean*. Taxonomic assignations were carried out using the *ecotag* command and the ArctBorBryo plant reference database. A unique taxon was assigned to each sequence with taxa corresponding to the last common ancestor node in the National Center for Biotechnology Information (NCBI) taxonomic tree. If several matches between the query sequence and the reference database were possible, the sequence was assigned to the taxon corresponding to the last common ancestor node of all the taxa in the NCBI taxonomic tree that best matched against the query sequence. A species name was accepted only if the identity score strictly equaled 1.00, a genus name in cases where the best match was ≥ 0.98, and a family name if the maximum identity was ≥ 0.95. Datasets were imported in Rstudio (R version 4.1.2) for further curation. Molecular Operational Taxonomic Units (MOTUs) labeled as PCR errors were filtered out. PCR replicate outliers (likely corresponding to non-functional PCR reactions) were also discarded. For this, we calculated the Euclidean distances of PCR replicates with their average (hereafter *dw*) and compared it against the distribution of pairwise dissimilarities between all average samples (hereafter *db*). Based on the expectation that PCR replicates from the same sample should be more similar than any two average samples (*dw* < *db*), we discarded PCR replicates lying outside the dissimilarity threshold defined as the intersection of *dw* and *db* distributions. This process was repeated iteratively until no more PCR replicates were removed from the dataset. If only a single PCR replicate per sample was left at the end, the sample was removed from the dataset. At the end of this procedure and in order to give equal weight to each replicate, remaining PCR replicates were averaged for each sample. Sequence reads abundance was normalised by dividing the number of reads for each MOTU by the total number of reads for each sample. MOTUs with relative abundance <1% in at least one sample as well as MOTUs with a best-identity match below 94% were discarded. Finally, PCR amplification success, tag jumps and potential cross-contaminations among samples were assessed by inspecting the detection levels and sequence reads abundance patterns of the synthetic standard sequences used as PCR positive controls. MOTUs were manually checked against the Svalbard flora checklist (http://svalbardflora.no) and taxonomic identifications refined whenever possible. MOTUs were grouped into eight functional categories based on Bråthen et al. (2007).

### Microbiome 16S amplicon sequencing

Library preparation for DNA sequencing was carried out as previously described (de Muinck et al. 2017), targeting the V4 region of the 16S rRNA gene with the 515f-805r primer pair. 2×300 bp pair-end sequencing was performed using the MiSeq platform at the Norwegian Sequencing Centre, Oslo. Sequence read demultiplexing was carried out using a custom routine developed at the Norwegian Sequencing Centre (https://github.com/nsc-norway/triple_index-demultiplexing). Further sequence data processing was performed using the Divisive Amplicon Denoising Algorithm as implemented in the *dada2* v1.16 R package (Callahan et al. 2016). Taxonomic classification of amplicon sequence variants (ASVs) was done using the SILVA v138.1 16S rRNA gene reference database (Quast et al. 2013). The read depth was adjusted through common scaling to the smallest reads depth (28 430 reads). Singletons as well as reads identified as either mitochondria or chloroplasts were removed from the data.

### Statistical analyses

All analyses were run with R (version 4.1.2). Diet diversity for each individual was calculated across MOTU’s using Hill’s numbers (Hill 1973; Keylock 2005) for q-value=1 corresponding to the exponential of Shannon’s entropy index (Spellerberg & Fedor 2003). Microbiome richness and diversity were estimated using Chao1 and the Shannon’s entropy indices. The inter-annual variation in major components of both the diet (proportion of grasses and proportion of *Salix*) and the gut microbiome (Bacteroidota or Firmicutes phyla) were arcsine transformed and analysed using a linear regression model. Year and valley were first fitted as categorical variables (degrees of freedom = 6 and 3, respectively). Linear trends over time (year as a linear variable) or in one of the environmental variables reflecting plant productivity (maximum NDVI or July temperature) were tested in competing models (after removing year as a categorical variable). The analysis of body mass in relation to the proportion of *Salix* in the diet was analysed using a generalized additive mixed model (function ‘*gamm’* in the *lme4* package) with year as random intercept, with the proportion of *Salix* in the diet, reproductive status (lactating, non-lactating), valley, and two-way interactions fitted as linear effects. Age was fitted with a spline function because of the non-linear relationship with body mass. Assessing the effects of valley, year, and the amount of *Salix* in the diet PERMANOVA was carried out using the ‘*adonis2*’ functions with 10 000 permutations and Bray-Curtis distances in the *vegan* version 2.5.6 package. We set up the “by=terms” setting, considering “valley” as the first term, “year” as the second term, and the amount of *Salix* in the diet as the third and last term, in order to account for the confounding effects of the year and valley. Partial redundancy analysis (RDA) was carried out using the *rda* function in the *vegan* package. The data matrix was transformed using the “*hellinger*” option in the *decostand* function in *vegan*. In the model formulation, “valley” and “year” were included as conditioning variables in order to focus only on the effects of the proportion of *Salix* in the reindeer diet. Computation of p-value for the *Salix* term in the RDA model was done using the *anova.cca* in *vegan* function and with 10 000 permutations. We carried out a non-metric multidimensional scaling (NMDS) analysis using the *metaMDS* function and Bray-Curtis distances in the *vegan* package using the default settings, and contour lines were added with the *ordisurf* function from the same package. Generalized additive models (GAMs) were computed with the *gam* function in the *mgcv* package (Wood 2011), using 5 degrees of freedom for the smooth terms in order to accommodate non-linear relationships. To separate any direct effect of *Salix* on body mass from a possible indirect effect via differences in the microbiome composition a piecewise structural equation model was fitted with function “*psem*” from the package *piecewiseSEM* (Lefcheck 2016).

## Results

### October diet

After bioinformatic analysis and data filtering, we retrieved 20 431 428 sequence reads from the samples passing all quality criteria. After removing data for calves and yearlings as well as failed samples, the final dataset comprised 97 individuals (Table S2). The Svalbard reindeer diet in October comprised 39 MOTUs from 14 different plant families. All plant MOTUs were identified at the family level, 33 (85%) at the genus level, and 13 MOTUs (33%) were identified at the species level (Table S3). The variation in the relative proportions across the eight functional groups is illustrated in Figure 1a. Diet diversity in October differed significantly between valleys (F_3,92_ = 6.59, p < 0.001) and tended to increase over time (F_1,92_ = 3.03, p = 0.085). Diet diversity increased significantly following warmer summers (July temperature, F _1,92_ = 4.02, p = 0.048; Figure 2a). The proportion of grass in the diet was generally low (mean = 0.045 ±0.068SD), and negatively correlated (p = 0.007) with the most common dietary item, the dwarf shrub *Salix polaris* (mean = 0.639 ±0.315SD), the only willow species on Svalbard. The proportion of *Salix* in diet drove the dietary diversity indices (Figure S2), with lowest diversity associated with both very high and very low proportions of *Salix*, while higher diversity occurred at intermediate *Salix* proportions and where grass was also relatively common (Figure S2). There were significant differences between valleys in the proportion of ingested *Salix* (F_3,87_ = 30.8, p < 0.001), from least in Eskerdalen (0.245), Sassendalen (0.468), Colesdalen (0.745), to most in Semmeldalen (0.915) (Figure S3). After accounting for valley differences, there was significant variation in the proportion of *Salix* in diet between years (F_6,87_ = 4.42, p < 0.001) but these showed no significant simple trend over time (p = 0.53). However, between year differences in *Salix* intake were associated positively with maximum NDVI (p = 0.009, Figure 2b), but not with July temperature (p = 0.4). While differences between valleys in the proportion of grass in the diet only tended towards significance (p = 0.068) this did vary significantly between years (F_6,87_ = 3.77, p < 0.002), and was associated positively with mean July temperature (p = 0.015, Figure 2c) but not with maximum NDVI (p = 0.71).

**Figure 1.**
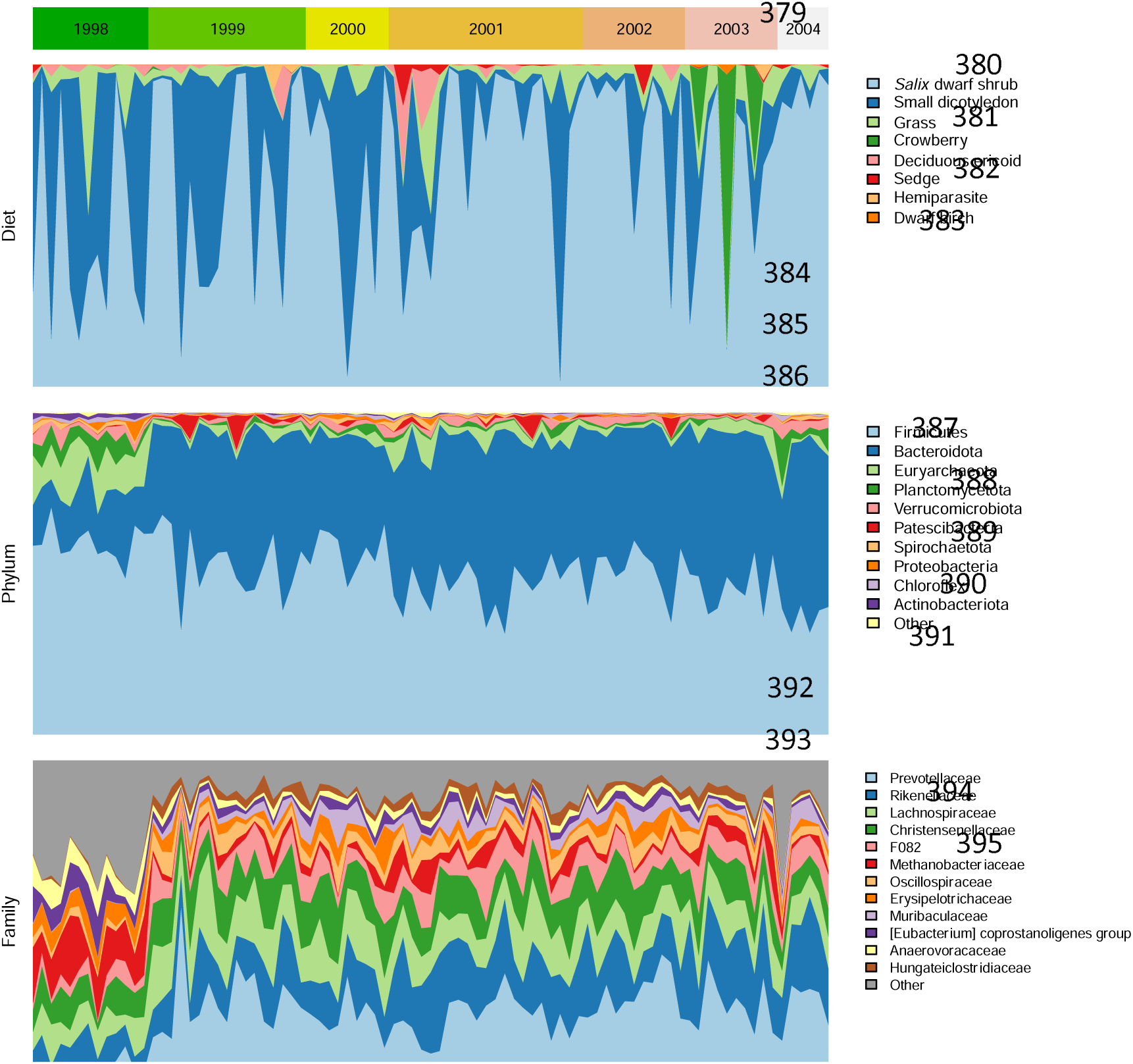
Temporal variation in the composition of the Svalbard reindeer diet (plant functional groups, upper panel) and the rumen microbiome (phylum and family level, lower two panels) over time. The width of the year bars is proportional to the sample size.

**Figure 2.**
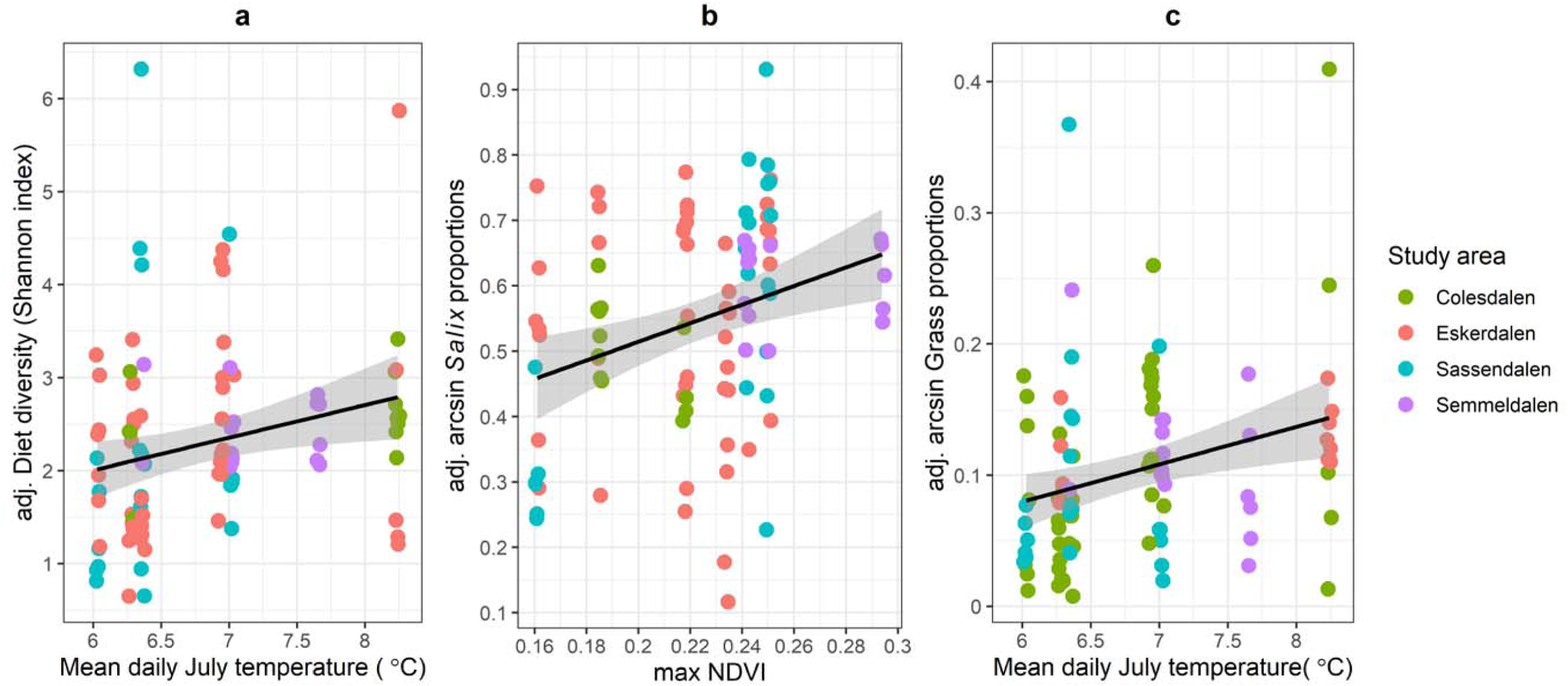
Diet diversity (Shannon index) of individuals plotted against mean July temperature (a), the arcsine proportion of *Salix* in the reindeer diet plotted against maximum NDVI (b) and the adjusted arcsine proportion of grass in the reindeer diet plotted against mean July temperature (c). The y-axis is adjusted for differences between valleys (coloured dots).

### Body mass and diet

After accounting for effects of valley and age, the proportion of *Salix* in diet had a positive effect on the October body mass, but only for lactating females (Figure 3a, Table S4). While the predicted increase in mass was as high as 7.6 kg from a *Salix*-free to an all-*Salix* diet, the equivalent estimate was 1.8 kg for non-lactating females, with the interaction close to significance (p = 0.055; Table S4). Despite differences in the average reindeer body mass between the valleys there was no significant interaction among lactating females between *Salix* proportion in diet, and valley (Figure 3b, all p > 0.25, Table S4).

**Figure 3.**
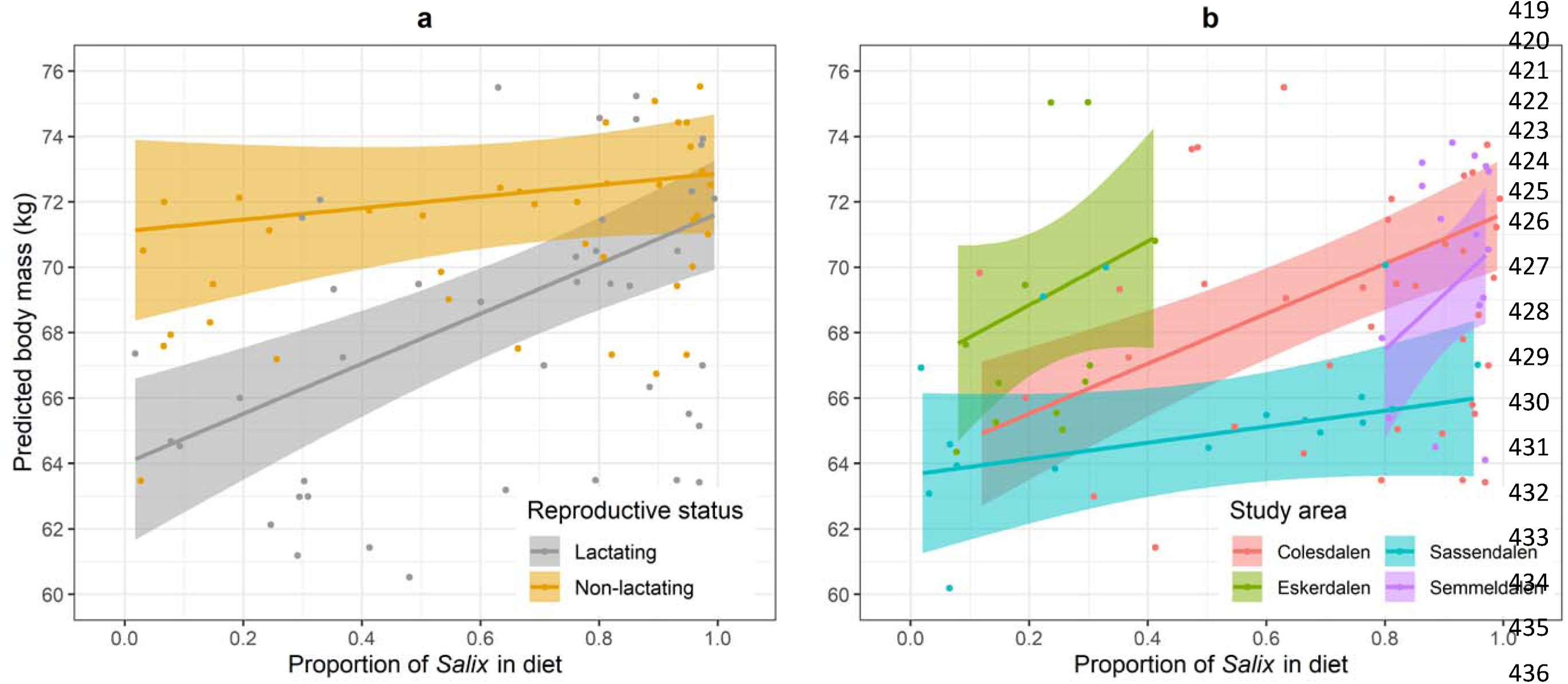
The relationship between the October body mass and the proportion of *Salix* dwarf shrub in the Svalbard reindeer diet for (a) lactating and non-lactating females, and for (b) lactating females from each of the four sampling valleys.

### The gut microbiome

For 87 of the 97 individuals used in the diet analysis, we also retrieved matching gut microbiome data. For the microbiome data, the total read number was 5 434 536 (mean: 62 466; SD: 13 831) matching 5 040 ASVs after quality filtering. We identified 24 unique bacterial phyla within the reindeer rumen microbiome with Bacteroidota (0.385 ±0.13SD) and Firmicutes (0.428 ±0.084SD) being the most abundant ones (Figure 1b). More than 60% of bacterial ASVs could be assigned to the family level with a total of 138 families identified (Figure 1c). Taxonomic classification accuracy of microbiome data is presented in Figure S4. Accounting for 80% of the reads, the proportion of Firmicutes and the proportion of Bacteriodota were highly negatively correlated across individuals (r = -0.729, p <0.0001). The proportion of Bacteriodota did not differ significantly between valleys (Wald Statistic = 2.21, d.f. = 3, p = 0.41, Figure S5) but increased over the course of the study (Wald Statistic = 12.21, d.f. = 1, p = 0.007, Figure 1). In contrast to Bacteriodota, Firmicutes differed between valleys (Eskerdalen = 0.379, Sassendalen = 0.382, Semmeldalen 0.445, Colesdalen = 0.457: Wald’s statistic = 19.96, d.f. = 3, P <0.001) and decreased significantly over the course of study (Wald Statistic = 21.97, d.f. =1, P = 0.001). Both the positive temporal trend in Bacteriodota and the negative trend in Firmicutes could be slightly better explained by variation in maximum NDVI (reduction in deltaAIC was -1.79 and -3.27, respectively, Figure 4). Despite the changes in Bacteriodota and Firmicutes the overall diversity of the microbiome did not change over the course of the study (Figure S6).

**Figure 4.**
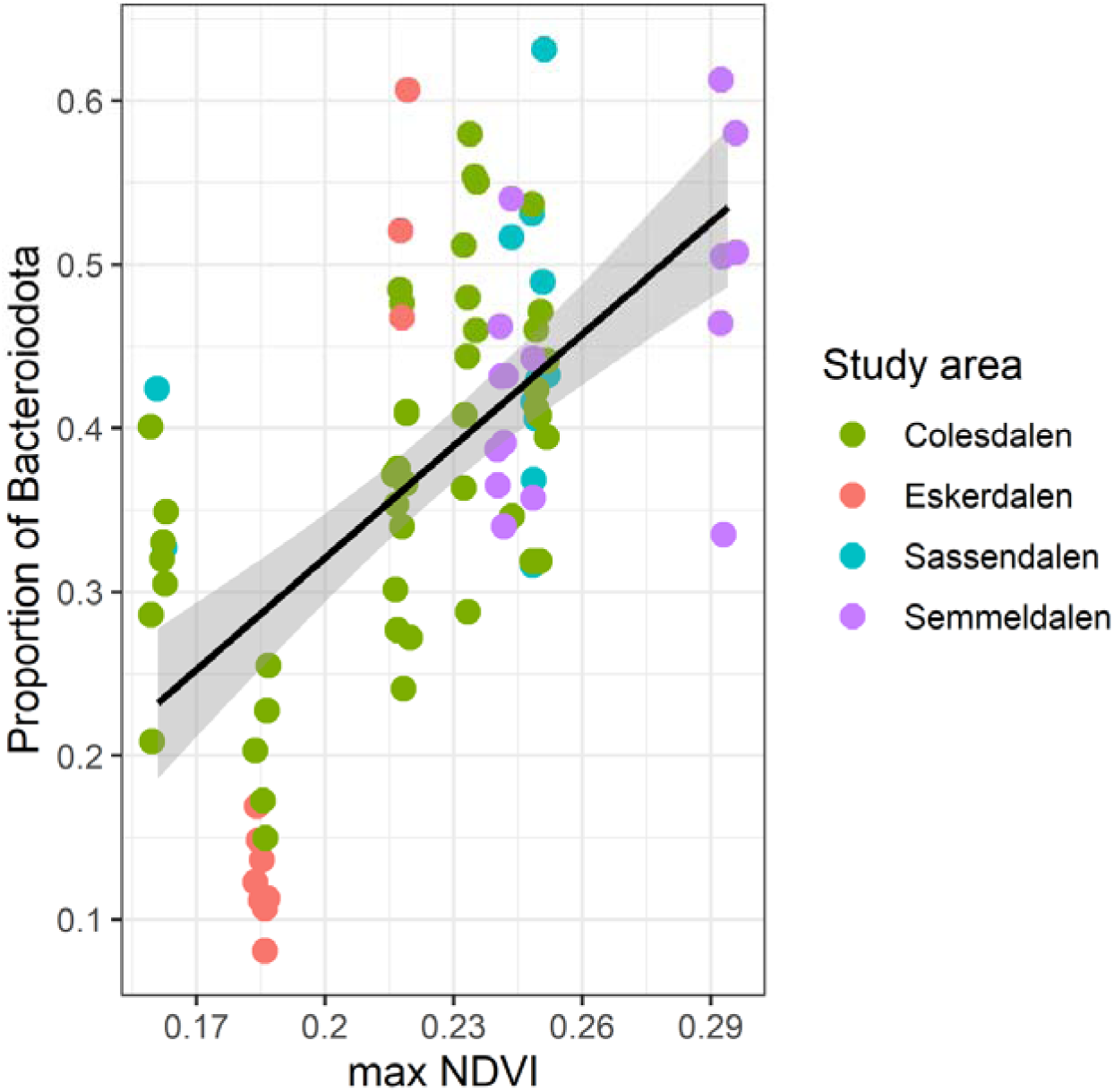
The proportion of Bacteroidota, the main bacterial phylum within the Svalbard reindeer rumen microbiome, plotted against maximum NDVI. The y-axis is adjusted for differences between valleys (coloured dots).

### Diet-microbiome interactions

At the individual level, the proportion of *Salix* in diet explained a significant amount of the variation in the gut microbiome composition (Figure 5, p = 0.012, PERMANOVA), after accounting for the potential confounding effects of the sampling valley and year. The effect of dietary *Salix* on the rumen microbiome was also confirmed with a separate modelling approach using partial redundancy analysis (p = 0.0015, PERMANOVA). Moreover, both Shannon entropy and Chao1 microbiome diversity measures were significantly positively correlated with the proportion of *Salix* in the diet (Figure 6). In addition, after controlling for the proportion of *Salix* in the diet, there was an independent negative effect of July temperature (Chao1 index: Wald statistic = 7.40, p = 0.033; Shannon entropy index Wald statistic = 9.21, P = 0.027). The piecewise structural equation model indicated that the contribution of the direct path effect of the proportion of *Salix* on body mass (partial correlation = 0.36 for Shannon; 0.41 for Chao1, after removing the effect of the significant correlation with the microbiome diversity) was greater than the indirect path effect through the microbiome diversity (0.29 x -0.26 = -0.08 for Shannon; 0.39 x -0.31 = -0.12 for Chao1) (Figure 7). Finally, after accounting for valley differences in diversity measures there was no simple relationship between diet diversity and microbiome diversity (Shannon’s entropy index: b = 0.011 ±0.027SE, Chao1 index: b = -0.04 ±8.8SE).

**Figure 5.**
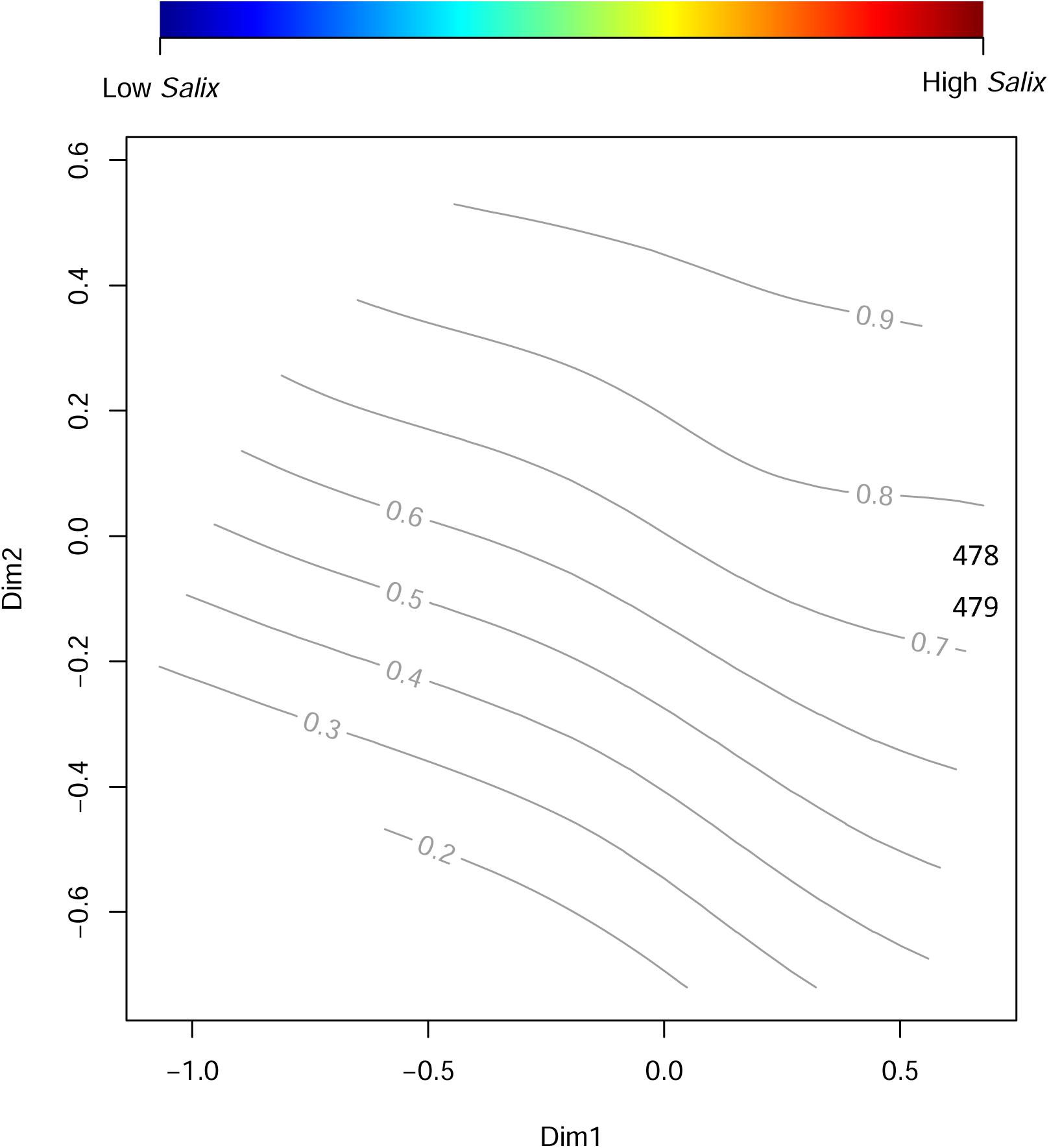
Non-metric multidimensional scaling of the Bray-Curtis distance matrix of the Svalbard reindeer rumen microbiome composition according to the proportion of *Salix* dwarf shrub (*Salix polaris*) in the reindeer diet. The effect is significant (p = 0.012, PERMANOVA).

**Figure 6.**
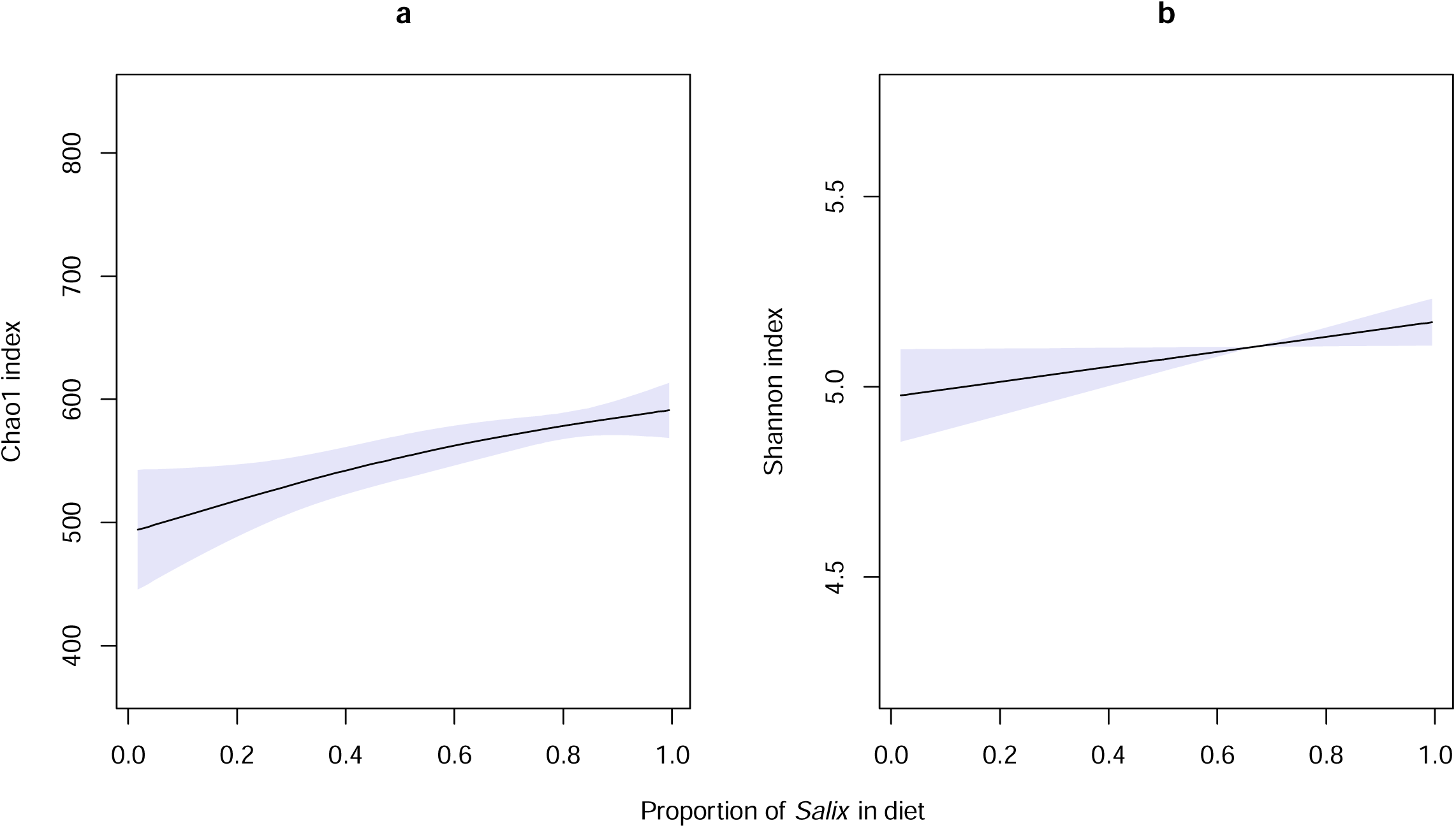
Generalised additive models showing the relationship between measures of microbiome diversity and the proportion of *Salix* in the diet of individual reindeer with the shaded bands indicating 95% confidence limits of the models. a) Chao1 index (p=0.007), and b) Shannon index (p=0.039).

**Figure 7.**
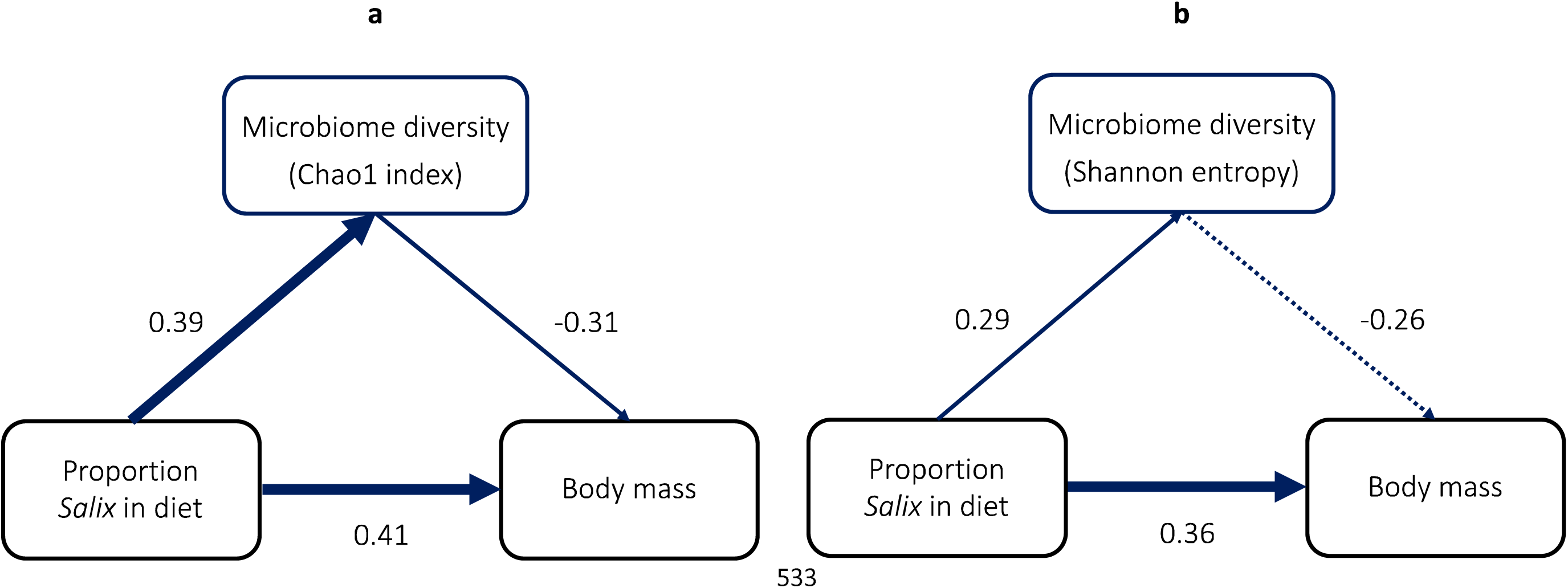
A piecewise structural equation model showing the direct and indirect (through microbiome) effect of *Salix* proportion in diet on age-specific body mass for lactating females. The values on the arrows are the standardized path coefficient and are effectively partial correlation coefficients, accounting for the correlation with the third variable. Arrow width is proportional to the strength of the coefficient.

## Discussion

An increase in plant biomass at a circumpolar scale is one of the major manifestations of climate warming in the Arctic tundra biome (Berner et al. 2020), potentially increasing carrying capacity for herbivore species. However, studies focusing on the impact of Arctic greening on large herbivores’ foraging, fitness and population consequences remain surprisingly rare. The sparse literature suggests both positive (Gagnon et al. 2020) and negative impacts on caribou (Fauchald et al. 2017). By combining a multi-year diet and gut microbiome DNA metabarcoding datasets for a keystone Arctic herbivore, we demonstrate a significant positive effect of the consumption of the major dwarf shrub on Svalbard, *Salix polaris*, on both autumn body mass and gut microbiome composition and diversity of individual Svalbard reindeer. Annual variation in the consumption of *Salix* was significantly, positively, related to the maximum NDVI, reflecting higher primary productivity. Thus, our study pinpoints an important mechanism mediating the ecological impact of climate warming on body mass, a trait influencing fertility and overwinter survival (Albon et al 2017), and a likely explanation for the growth of Svalbard reindeer populations (Le Moullec et al 2019).

Although diet diversity increased over time, the major finding of our study is that the higher intake of a single plant species – *Salix polaris* – leads to a higher autumn body mass, an effect that was particularly pronounced for females bearing the cost of lactation over the summer. Between-year variation in the proportion of grass in the diet, but not *Salix*, was significantly related to July temperature as expected based on the temperature-driven productivity measurements across plant functional types, recorded in the study area since 1998 (van der Wal & Stien 2014). However, *Salix* growth was significantly related to NDVI. Since we now know that annual variation in NDVI over a decadal time scale in our study area is strongly influenced by summer temperature (Karlsen et al. 2018), the lack of a direct correlation with July temperature may reflect that our seven-year study is a relatively short times series for consistently detecting trends. For example, 1998, the first year was the warmest of the seven we had dietary measures but had the second lowest maximum NDVI. Furthermore, the annual growth of *Salix* is not only dependent on July temperature but also negatively impacted by rain-on-snow events in the previous winter (Le Moullec et al 2020). These rain-on-snow events are stochastic and also tend to be localised, which could well contribute to differences in annual *Salix* growth, and hence availability, between valleys. Finally, while above-ground grass shoots are produced annually, shrub biomass accumulates over several years with a potential lagged positive effect of previous warm summers. This may contribute to explain why dietary proportion of *Salix* was best explained by NDVI (capturing shrubification over time) while proportion of grass was better explained by the current year’s July temperature.

*Salix polaris* is a high-quality resource for Svalbard reindeer (Bjørkvoll et al. 2009) and likely to play an important nutritional role in October when vegetation has senesced, and most plant nutrients are relocated to the underground parts. A significant proportion of the biomass of dwarf shrubs is buried below-ground but shallowly enough in the layer of soil to be accessible to herbivores after the above-ground biomass has senesced (Iversen et al. 2015; Le Moullec et al. 2018). This could explain discrepancies with earlier dietary studies based on micro-histology, showing that the Svalbard reindeer autumn diet tends to be dominated by grasses rather than *Salix*, in both rumen (Bjørkvoll et al. 2009) and faeces (Bjune 2000). On the one hand, with micro-histology grasses often tend to be overrepresented in faecal samples, at the expense of more digestible taxa such as forbs (Vavra & Holechek 1980; McInnis et al. 1983; Parker & Bernard 2006). On the other hand, micro-histology relies on the visual detection and identification of plant cell wall structures and stomata, which are absent from tissues such as roots.

While there was a strong positive effect of *Salix* intake in October on body mass, there was wide range of variation in *Salix* proportions in diet between individuals within years and valleys, suggesting this resource might not be equally accessible or preferred by all reindeer. Furthermore, since the proportion of *Salix* in rumen samples likely corresponds to foraging over one or few days (Barboza et al. 2006; Steuer et al. 2011; Picard et al. 2015), the ‘snapshot’ might not reflect the overall foraging strategy or preferences of individuals. Thus, the question is to what extent *Salix* is the ultimate driver of higher body mass or is representative of a more proximate overall foraging strategy.

Contrary to our expectations, and despite a significant relationship between *Salix* intake and gut microbiome composition and diversity, we did not detect a positive effect on reindeer body mass via the gut microbiome but a negative effect, which in the case of the Chao1 index was significant. The gut microbiome is a relatively new element in ecological studies, and our understanding of the role and relative importance of the gut bacteria on their mammal host’s ecology and fitness remains limited outside the human model (Suzuki 2017; McKenney et al. 2018). However, our results confirm the general understanding that dietary choice and gut microbiome composition are tightly linked (Muegge et al. 2011; Kartzinel et al. 2019; Baniel et al. 2021) as the characteristics of the dietary substrate determine which bacterial metabolic profiles will dominate the microbiome community. In our case, reindeer with a high proportion of *Salix* proportion in the rumen displayed more similar gut microbiomes and higher microbiome diversity, suggesting direct link with the foraging of this plant species by the reindeer. The higher microbiome diversity could be explained by the ingestion of the high-fiber woody *Salix* tissues requiring the recruitment of the gut microbiome community to aid with digestion (McKenney et al. 2018). However, the negative association between microbiome and reindeer body mass is more difficult to explain. Wild ungulates are constrained in their dietary choice by resource availability, often leading to foraging on plants of low nutritional quality or with high amounts of toxic secondary metabolites (Foley & Moore 2005; Iason 2007; Windels & Hewitt 2011). This most likely explains the general lack of a simple relationship between diet and gut microbiome diversity in large mammalian herbivores (Kartzinel et al. 2019), including high Arctic species (Prewer et al. 2023), as a dynamic trade-off might exist between meeting nutritional demands and the capacity to detoxify a wide range of plant defense compounds (Felton et al. 2018). Large mammalians gut microbiome plays an essential role in both functions (McKenney et al. 2018; Dearing & Weinstein 2022). This could explain the negative association between microbiome and reindeer body mass here as there might be a fine balance between gaining nutritional benefits from *Salix* ingestion in October and the capacity of the gut microbiome to detoxify deterrent secondary compounds. More recent, independent measurements from the same study area carried out in 2022 do show that shrubs such as *Salix* and *Saxifraga* exhibit among the highest concentrations of phenolics compared to forbs and grasses, and that these concentrations remain high even during the autumn (Eikeland 2023).

On the other hand, diverse gut microbiomes have been also associated with higher exploratory behaviour, behavioural flexibility, and the associated foraging success (Davidson et al. 2018; Florkowski & Yorzinski 2023), suggesting that the positive effect of gut microbiome diversity could be confounded with foraging, rather than acting independently on the reindeer body mass. For example, a recent study on the mountain goat (*Oreamnos americanus*) in British Columbia and the white-tailed deer (*Odocoileus virginianus*) in Ontario found correlations between the gut microbiome composition and home range size (Wolf et al. 2021). Home range size could be an important indicator of activity levels, as well as the energetic cost of foraging, suggesting the beneficial impact of altered microbiomes in response to dietary change could also be expressed through impact on less intuitive fitness traits.

Interestingly, we also observe a temporal trend in increasing Bacteroidota proportions, and commensurate decline in Firmicutes, in the reindeer rumen over time, correlating positively and negatively, respectively, with NDVI. The significance of this apparent temperature-induced change is unknown. Nonetheless, there is an increasing amount of evidence that changes in the mean environmental temperature can modify gut microbiome composition and diversity, independently from host diet and behaviour (Sepulveda & Moeller 2020; Li et al. 2022; Williams et al. 2023), but how such changes then impact on host fitness remains understudied. However, considering the seemingly tight link between diet and gut microbiome composition in our study, our findings open the question of whether the gradual change in microbiome composition over time could be considered as indicative of a general shift towards improved diet quality in the Svalbard reindeer, which would be otherwise difficult to capture with measures such as DNA metabarcoding (i.e., short-term diet), or NDVI (i.e., coarse measure of plant greenness, not always reflecting resource availability, especially at local scales).

## Conclusion

Our findings provide a mechanistic understanding of how a keystone herbivore species has benefitted from climate warming. Diet diversity increased over the course of our seven-year study, driven by a higher ingestion of *Salix polaris* following summers of higher plant productivity (as indexed by the maximum NDVI), with grasses also more common following warmer summers. Among individuals, a higher proportion of *Salix* in the autumn diet led to higher body mass, which in turn is known to improve reproduction and overwinter survival. Interestingly, there was only a relatively weak, and unexpectedly negative effect of microbiome diversity on body mass, independent of the positive effect of *Salix*. These results suggest that, while overall microbiome diversity is resilient to substantial changes in diet, major phyla may still exhibit marked changes, potentially providing resilience for the host to dietary changes induced by climate warming. Our study highlights the value of combining fine-grained DNA metabarcoding data to unravel the effect of diet−microbiome linkages on the fitness and population growth of wild-ranging ungulates, with implications relevant for herbivore populations challenged by climate warming.

## Acknowledgements

We warmly thank the numerous students and researchers assisting with the Svalbard field campaigns. We thank the Governor of Svalbard for permission to undertake the research and technical staff at the University Centre in Svalbard (UNIS) and the Norwegian Polar Institute for supporting field campaigns and providing access to lab facilities. We are also particularly grateful to Éric Coissac for his overall scientific contribution as well as assistance with data analysis and interpretation. This work was supported by the Research Council of Norway through projects 257642 (“REININ: Reindeer interactions from plants and birds to humans: balancing the odds of climate change”) and 315454 (“PRISM: Understanding climate change impacts in an Arctic ecosystem: an integrated approach through the prism of Svalbard reindeer”). SK was also partially funded by the EU Horizon 2020 Programme for Research and Innovation through Action Number 869471 (CHARTER: Drivers and feedbacks of changes in arctic terrestrial biodiversity, PI: Bruce Forbes).

## Author’s contributions

All authors contributed to ideas and the conception of the study. R. Justin Irvine, Rolf Langvatn, Steve D. Albon and Leif Egil Loe are part of the Svalbard reindeer monitoring project, responsible for data collection and management. Stefaniya Kamenova carried out DNA metabarcoding diet analysis with input from Galina Gusarova, while Eric Jacques de Muinck and Pål Trosvik carried out gut microbiome analysis. Pål Trosvik, Steve D. Albon and Leif Egil Loe carried out statistical analyses. Stefaniya Kamenova wrote the first draft with input from Steve D. Albon, and all authors contributed to the final version of the manuscript.

## Data availability statement

Raw sequencing datasets are available at the Sequence Read Archive database under accession numbers PRJNA1052483 (DNA metabarcoding diet data) and PRJNA1044740 (16S amplicon microbiome data). Metadata and scripts will be made available upon publication.

## Permits

Reindeer sample collection met all guidelines of the Research Council of Norway and were collected with the permission of the Governor of Svalbard (Sysselmannen).

