## Supplementary material for "Arctic greening drives changes in the diet and gut microbiome of a resident herbivore with consequences for fitness": Table S1

**Table S1.** Details about the six synthetic DNA standards used as a positive control mock community, amplified with the *Sper01* seed plant primers as part of the DNA metabarcoding diet analysis of the Svalbard reindeer.

| Synthetic standard | Dilution | Reference sequence | Standard length (bp) | Standard GC content (%) |
| --- | --- | --- | --- | --- |
| SS_01_GH | 1 | taagtctcgcactagttgtgacctaacgaatagagaattctataagacgtgttgtcccat | 60 | 40 |
| SS_02_GH | 1/2 | gtgtatggtatatttgaataatattaaatagaatttaatcaatctttacatcgcttaata | 60 | 20 |
| SS_03_GH | 1/4 | cacaatgctcggtaactagaagcatttgta | 30 | 40 |
| SS_04_GH | 1/8 | attgaatgaaaagattattcgatatagaat | 30 | 20 |
| SS_05_GH | 1/16 | agaacgctagaatctaagatggggggggggatgagtaagatatttatcagtaacatatga | 60 | 40 |
| SS_06_GH | 1/32 | atttttgtaactcattaacaattttttttttgatgtatcataagtactaaactagttact | 60 | 20 |

**Table S2.** Number of individual Svalbard reindeer rumen samples per year and locality retained in the final diet-gut microbiome matched dataset.

|  | Colesdalen | Eskerdalen | Sassendalen | Semmeldalen |
| --- | --- | --- | --- | --- |
| 1998 | 7 | 8 | 0 | 0 |
| 1999 | 13 | 5 | 0 | 0 |
| 2000 | 7 | 0 | 2 | 0 |
| 2001 | 10 | 0 | 10 | 2 |
| 2002 | 1 | 0 | 1 | 9 |
| 2003 | 10 | 0 | 0 | 0 |
| 2004 | 0 | 0 | 0 | 8 |

**Table S3.** Overall plant diet composition of the Svalbard reindeer assessed with DNA metabarcoding from 97 rumen content samples, collected in October between 1998 and 2004.

| **Best identity match** | **Family** | **Genus** | **Species** | **Scientific name** | **Assumed taxon** | **Functional category** |
| --- | --- | --- | --- | --- | --- | --- |
| 1 | Salicaceae | NA | NA | Saliceae | Salix sp. | Salix dwarf shrub |
| 1 | Saxifragaceae | *Saxifraga* | *Saxifraga oppositifolia* | *Saxifraga oppositifolia* | *Saxifraga oppositifolia* | Small dicotyledon |
| 1 | Poaceae | NA | NA | Pooideae | Poaceae | Grass |
| 1 | Poaceae | *Festuca* | NA | *Festuca* | *Festuca* sp. | Grass |
| 1 | Polygonaceae | *Bistorta* | *Bistorta vivipara* | *Bistorta vivipara* | *Bistorta vivipara* | Small dicotyledon |
| 0.98 | Saxifragaceae | *Saxifraga* | *Saxifraga oppositifolia* | *Saxifraga oppositifolia* | *Saxifraga oppositifolia* | Small dicotyledon |
| 1 | Rosaceae | *Dryas* | NA | *Dryas* | *Dryas octopetala* | Deciduous ericoid |
| 1 | Ericaceae | *Empetrum* | NA | *Empetrum* | *Empetrum nigrum* | Crowberry |
| 1 | Brassicaceae | *Draba* | NA | *Draba* | *Draba* sp. | Small dicotyledon |
| 1 | Saxifragaceae | *Saxifraga* | NA | *Saxifraga* | *Saxifraga* sp. | Small dicotyledon |
| 1 | Saxifragaceae | *Saxifraga* | *Saxifraga cespitosa* | *Saxifraga cespitosa* | *Saxifraga cespitosa* | Small dicotyledon |
| 1 | Polygonaceae | *Oxyria* | *Oxyria digyna* | *Oxyria digyna* | *Oxyria digyna* | Small dicotyledon |
| 1 | Poaceae | NA | NA | Poinae | Poaceae | Grass |
| 1 | Betulaceae | *Betula* | NA | *Betula* | *Betula nana* | Dwarf birch |
| 0.96 | Saxifragaceae | *Saxifraga* | NA | *Saxifraga* | *Saxifraga* sp. | Small dicotyledon |
| 1 | Poaceae | NA | NA | Poeae | Poaceae | Grass |
| 1 | Caryophyllaceae | *Cerastium* | NA | *Cerastium* | *Cerastium* sp. | Small dicotyledon |
| 0.98 | Saxifragaceae | *Saxifraga* | NA | *Saxifraga* | *Saxifraga* sp. | Small dicotyledon |
| 1 | Juncaceae | *Luzula* | NA | *Luzula* | *Luzula* sp. | Sedge |
| 0.98 | Salicaceae | NA | NA | Saliceae | *Salix* sp. | Salix dwarf shrub |
| 1 | Ranunculaceae | *Ranunculus* | Ranunculus pygmaeus | *Ranunculus pygmaeus* | *Ranunculus pygmaeus* | Small dicotyledon |
| 1 | Orobanchaceae | *Pedicularis* | NA | *Pedicularis* | *Pedicularis* sp. | Hemiparasite |
| 1 | Poaceae | NA | NA | Poeae | Poaceae | Grass |
| 1 | Papaveraceae | *Papaver* | NA | *Papaver* | *Papaver* sp. | Small dicotyledon |
| 1 | Caryophyllaceae | *Minuartia* | *Minuartia biflora* | *Minuartia biflora* | *Minuartia biflora* | Small dicotyledon |
| 1 | Ericaceae | *Cassiope* | *Cassiope tetragona* | *Cassiope tetragona* | *Cassiope tetragona* | Deciduous ericoid |
| 0.98 | Ericaceae | *Cassiope* | *Cassiope tetragona* | *Cassiope tetragona* | *Cassiope tetragona* | Deciduous ericoid |
| 0.97 | Ranunculaceae | *Ranunculus* | *Ranunculus pygmaeus* | *Ranunculus pygmaeus* | *Ranunculus pygmaeus* | Small dicotyledon |
| 0.98 | Rosaceae | *Dryas* | NA | *Dryas* | *Dryas octopetala* | Deciduous ericoid |
| 1 | Orobanchaceae | *Pedicularis* | NA | *Pedicularis* | *Pedicularis* sp. | Hemiparasite |
| 1 | Poaceae | *Puccinellia* | NA | *Puccinellia* | *Puccinellia* sp. | Grass |
| 1 | Cyperaceae | *Carex* | NA | *Carex* | *Carex* sp. | Sedge |
| 0.96 | Ericaceae | *Cassiope* | *Cassiope tetragona* | *Cassiope tetragona* | *Cassiope tetragona* | Deciduous ericoid |
| 1 | Cyperaceae | *Eriophorum* | NA | *Eriophorum* | Eriophorum sp. | Sedge |
| 1 | Saxifragaceae | *Saxifraga* | *Saxifraga hirculus* | *Saxifraga hirculus* | *Saxifraga hirculus* | Small dicotyledon |
| 1 | Saxifragaceae | *Saxifraga* | NA | *Saxifraga* | *Saxifraga* sp. | Small dicotyledon |
| 1 | Orobanchaceae | *Pedicularis* | *Pedicularis lanata* | *Pedicularis lanata* | *Pedicularis lanata* | Hemiparasite |
| 1 | Rosaceae | *Potentilla* | NA | *Potentilla* | *Potentilla* sp. | Small dicotyledon |
| 0.98 | Saxifragaceae | *Saxifraga* | NA | *Saxifraga* | *Saxifraga* sp. | Small dicotyledon |

**Table S4.** A generalised additive mixed model explaining variation in body mass of female Svalbard reindeer as a function of study area, proportion of *Salix* in the diet and lactation status, including two-way interactions. Age is accounted for in a separate non-linear spline term (estimated degrees of freedom = 4.4; p < 0.001) while year is fitted as a random intercept (with SD = 1.96).

|  | **Value** | **SE** | **t** | **p** |
| --- | --- | --- | --- | --- |
| Intercept | 62.4 | 2.5 | 24.93 | <0.001 |
| Eskerdalen vs Colesdalen | 2.9 | 4.1 | 0.70 | 0.487 |
| Sassendalen vs Colesdalen | -0.3 | 3.0 | -0.11 | 0.910 |
| Semmeldalen vs Colesdalen | -10.1 | 17.6 | -0.57 | 0.569 |
| Salix | 7.6 | 3.1 | 2.47 | 0.016 |
| Lactating (no vs yes) | 7.1 | 2.1 | 3.34 | 0.001 |
| Salix vs Lactating (no vs yes) | -5.9 | 3.0 | -1.94 | 0.055 |
| Eskerdalen x Salix | 2.1 | 14.3 | 0.15 | 0.883 |
| Sassendalen x Salix | -5.2 | 4.5 | -1.15 | 0.252 |
| Semmeldalen x Salix | 9.3 | 19.2 | 0.49 | 0.629 |

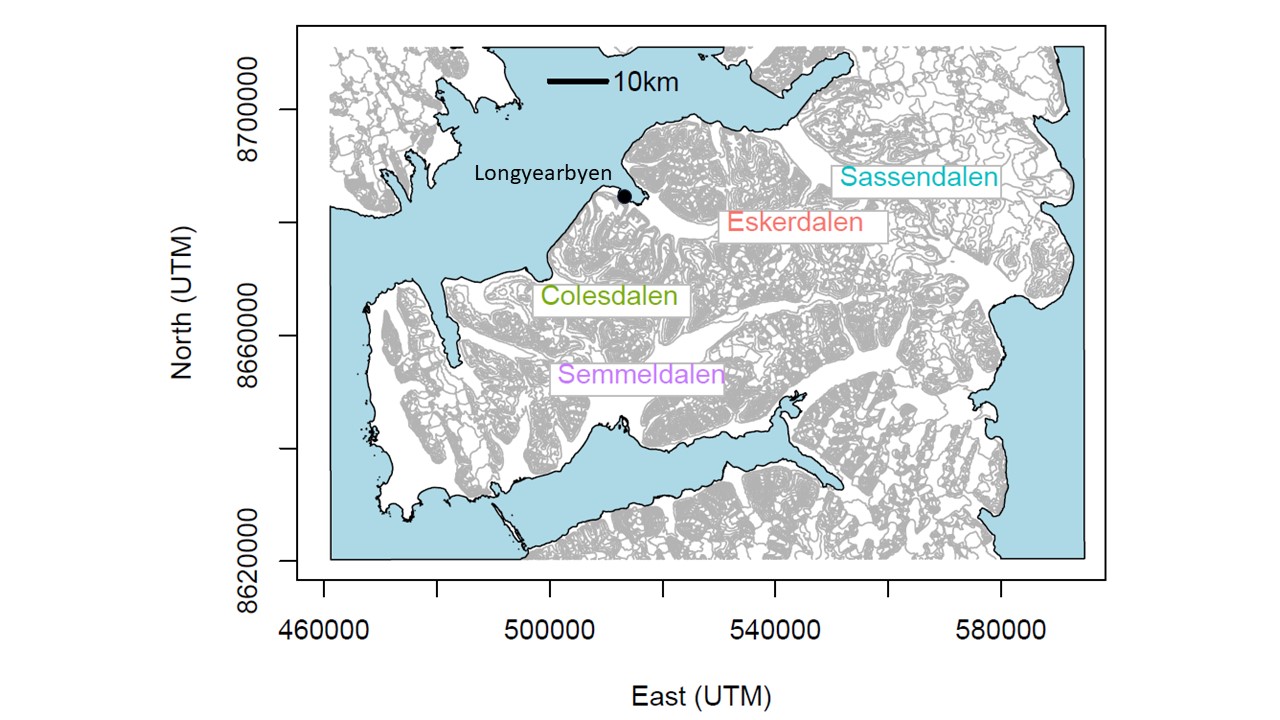

**Figure S1.** Map of the study area indicating the locations of the Colesdalen, Semmeldalen, Eskerdalen and Sassendalen valleys, and the main settlement Longyearbyen.

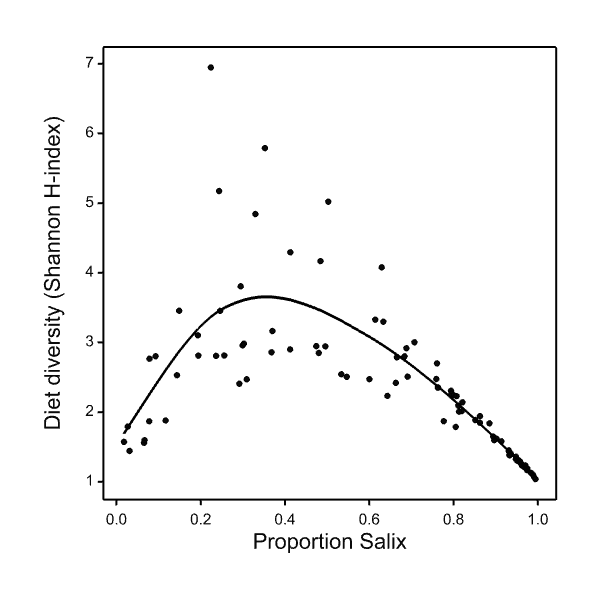

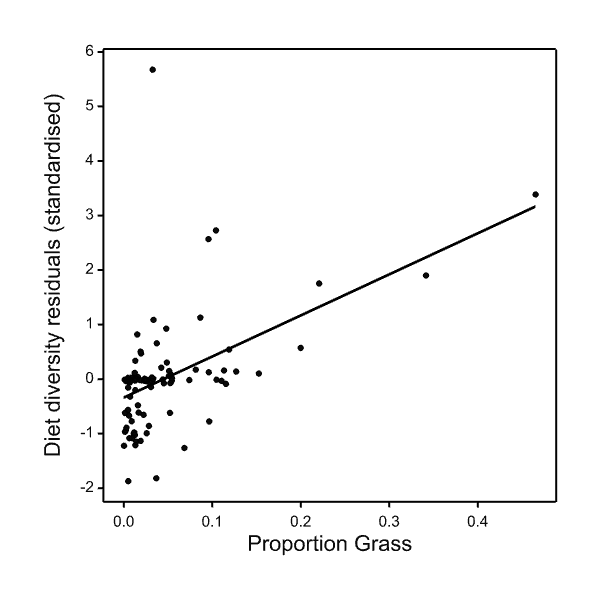

**Figure S2.** The relationship between diet diversity and proportion *Salix* in the diet (left-handed panel), and the standardised residual from the left figure plotted against the proportion grass in the diet (right-handed panel).

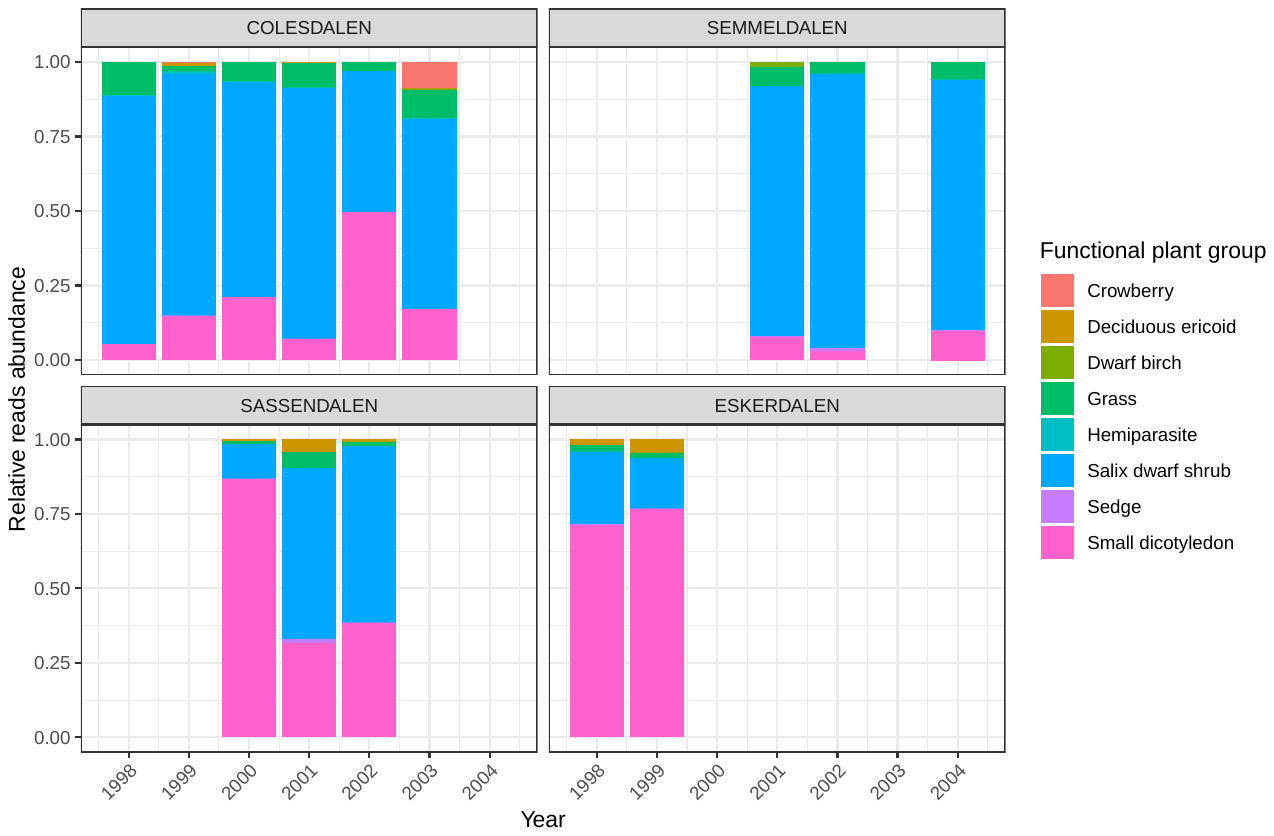

**Figure S3.** Variation over time of the relative proportion of the different plant functional groups, detected in the October diet of the Svalbard reindeer for each year and sampling location.

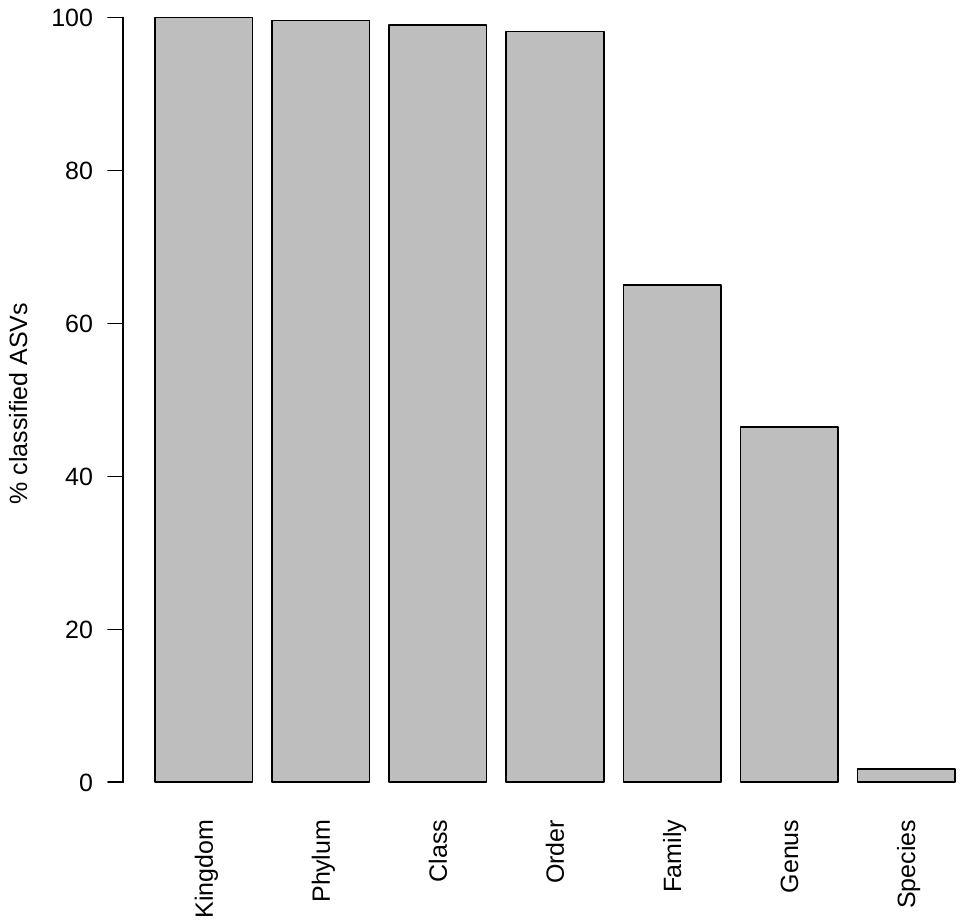

**Figure S4.** Percentage of Amplicon Sequence Variants (ASVs) classified to the indicated taxonomic level with >50% probability for the Svalbard reindeer rumen microbiome dataset.

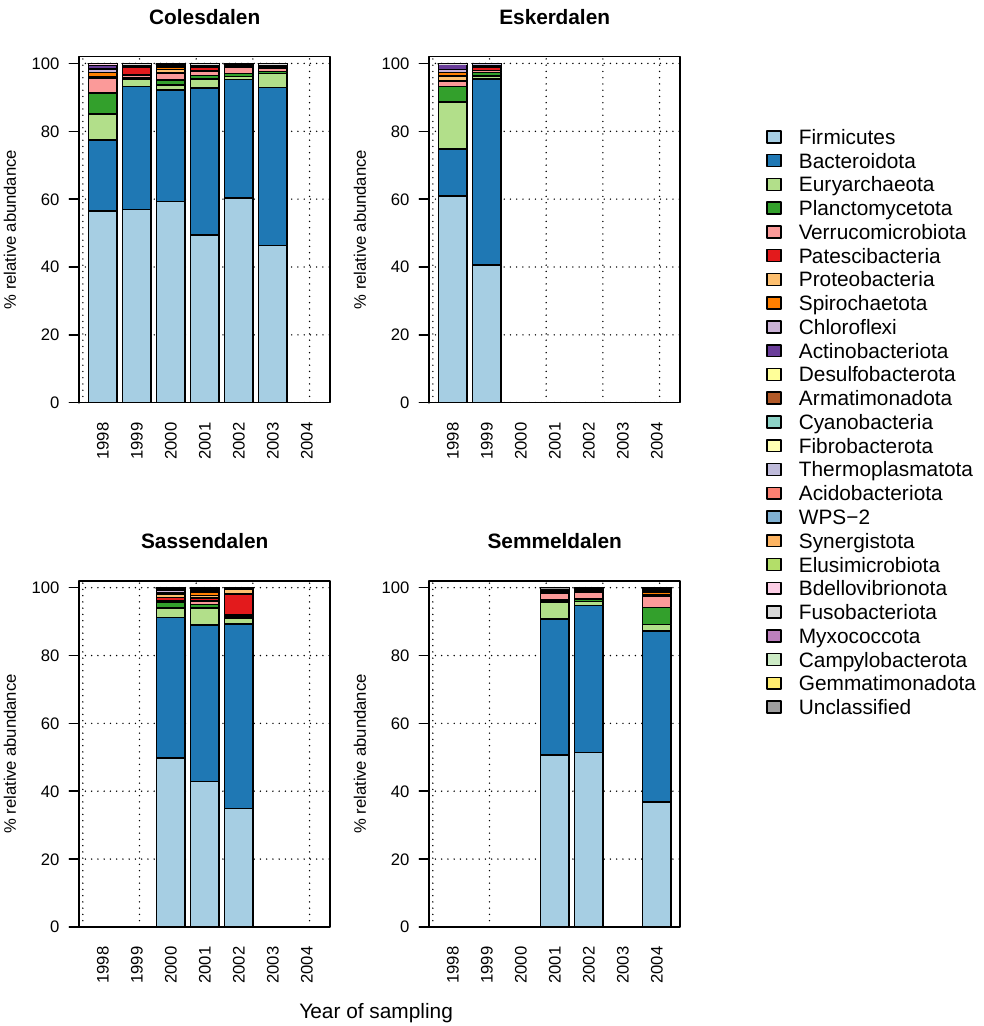

**Figure S5.** Rumen microbiome composition of the Svalbard reindeer, sampled in October in across four locations between 1998 and 2004.

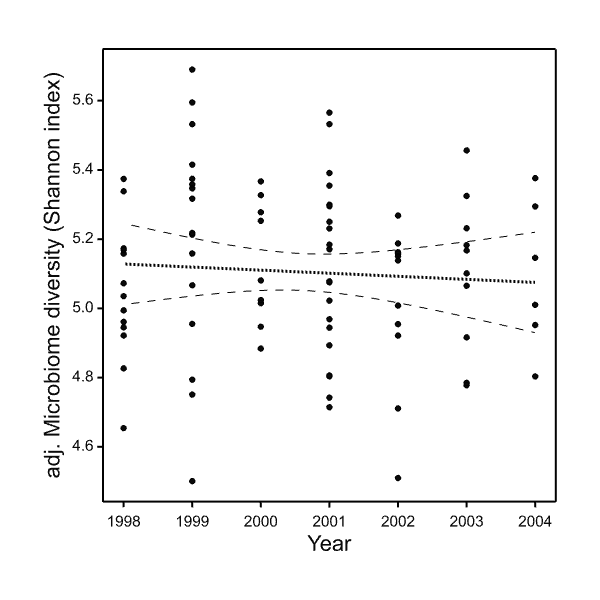

**Figure S6.** Microbiome diversity (Shannon entropy index) adjusted for valley differences and plotted against year.
